# Alterations of PINK1-PRKN signaling in mice during normal aging

**DOI:** 10.1101/2024.04.29.591753

**Authors:** Zahra Baninameh, Jens O. Watzlawik, Xu Hou, Tyrique Richardson, Nicholas W. Kurchaba, Tingxiang Yan, Damian N. Di Florio, DeLisa Fairweather, Lu Kang, Justin H. Nguyen, Takahisa Kanekiyo, Dennis W. Dickson, Sachiko Noda, Shigeto Sato, Nobutaka Hattori, Matthew S. Goldberg, Ian G. Ganley, Kelly L. Stauch, Fabienne C. Fiesel, Wolfdieter Springer

## Abstract

The ubiquitin kinase-ligase pair PINK1-PRKN identifies and selectively marks damaged mitochondria for elimination via the autophagy-lysosome system (mitophagy). While this cytoprotective pathway has been extensively studied *in vitro* upon acute and complete depolarization of mitochondria, the significance of PINK1-PRKN mitophagy *in vivo* is less well established. Here we used a novel approach to study PINK1-PRKN signaling in different energetically demanding tissues of mice during normal aging. We demonstrate a generally increased expression of both genes and enhanced enzymatic activity with aging across tissue types. Collectively our data suggest a distinct regulation of PINK1-PRKN signaling under basal conditions with the most pronounced activation and flux of the pathway in mouse heart compared to brain or skeletal muscle. Our biochemical analyses complement existing mitophagy reporter readouts and provide an important baseline assessment *in vivo,* setting the stage for further investigations of the PINK1-PRKN pathway during stress and in relevant disease conditions.

## INTRODUCTION

PINK1-PRKN directed mitophagy has emerged as a critical mitochondrial quality control pathway with likely far-reaching implications and relevance to stress, aging, and disease [1]. When mitochondria are damaged, the kinase PINK1 accumulates locally on the outer mitochondrial membrane where it phosphorylates ubiquitin (Ub) at serine-65 (p-S65-Ub) [2–4]. This signal acts as a specific receptor and allosteric activator of the E3 Ub ligase PRKN, which is recruited from the cytosol and structurally de-repressed [5–7]. PINK1 also phosphorylates a conserved serine-65 residue in the Ub-like domain of PRKN to fully activate and unleash the E3 Ub ligase [8–11]. PINK1 and PRKN then engage in a positive feedback loop to decorate damaged mitochondria with p-S65-Ub chains. These serve as a specific mitophagy label facilitating their autophagic delivery and elimination in lysosomes.

This process acts to maintain overall health and function of the mitochondrial network and thus is thought to be broadly cytoprotective. The importance of the pathway is further underscored as complete loss of PINK1 or PRKN function are the most common causes of early-onset Parkinson’s disease (PD) [12,13]. While PINK1-PRKN signaling seems particularly important for dopaminergic neurons that degenerate in PD [1,14,15], both enzymes are widely expressed in a variety of tissues and cell types. Alterations in PINK1-PRKN mitophagy are thought to play a role in many human age-related disorders including Alzheimer’s disease, cardiovascular disease, and myopathies and a prominent accumulation of p-S65-Ub is commonly seen in aging and disease [16–21].

PINK1 and PRKN have been extensively studied in cell culture often under overexpression conditions and upon acute and massive mitochondrial depolarization, but the relevance of PINK1-PRKN mitophagy is not well-established *in vivo* and debated [22]. In contrast to Drosophila, *pink1*^-/-^ or *prkn*^-/-^ mouse models only have subtle phenotypes and no change in overall mitophagy rates has been observed under basal conditions (reviewed in [23]). However, consistent with the idea of a stress-activated pathway, infections, exhaustive exercise, advanced aging, and proteotoxic conditions all seem to increase PINK1-PRKN activity in mice or aggravate phenotypes in their absence [17,24–27].

Recently, we demonstrated that PINK1-PRKN signaling is indeed active in different human cell types and in rodent brain under basal conditions, albeit at extremely low levels thus requiring sensitive detection methods [28,29]. Here we expanded our analyses and determined the basal activity of the PINK1-PRKN pathway in brain, heart, and skeletal muscle of young, middle-aged, or old wild-type (WT) compared to *pink1*^-/-^ or *prkn*^-/-^ mice. Our results highlight similarities and differences between tissues with regards to the extent and regulation of PINK1-PRKN signaling during normal aging which contribute to a better understanding of this pathway on the organismal level.

## RESULTS

### PINK1-PRKN signaling in brain, skeletal muscle, and heart of young mice

To determine PINK1-PRKN signaling in different organs, we first compared WT to age-matched *pink1*^-/-^ and *prkn*^-/-^ mice. Brain, skeletal muscle, and heart from young (3-month-old) animals were obtained and RNA and protein extracted. Quantitative real-time polymerase chain reaction (RT-PCR) confirmed the genotypes but there were no differences between expression of *Pink1* in *prkn*^-/-^ mice or *Prkn* in *pink1*^-/-^ mice in any of the tissues compared to WT (**Fig. S1A-B)**.

Due to lack of robust and reliable methods to measure mouse PINK1 protein, we next focused on PRKN and determined protein levels by western blot (**Fig. 1A**). PRKN protein was absent in the different organs from *prkn*^-/-^ mice, as expected. We further observed a slight but significant increase (∼1.3-fold) in PRKN protein in brain from *pink1*^-/-^ animals compared to WT, consistent with our recent report linking PINK1-mediated activation and turnover of PRKN [29]. While there was no such change noted in skeletal muscle, we observed an even greater increase in (inactive) PRKN protein (∼2.5-fold) in *pink1*^-/-^ hearts pointing to higher activation and turnover of PRKN in this organ.

**Figure 1.**
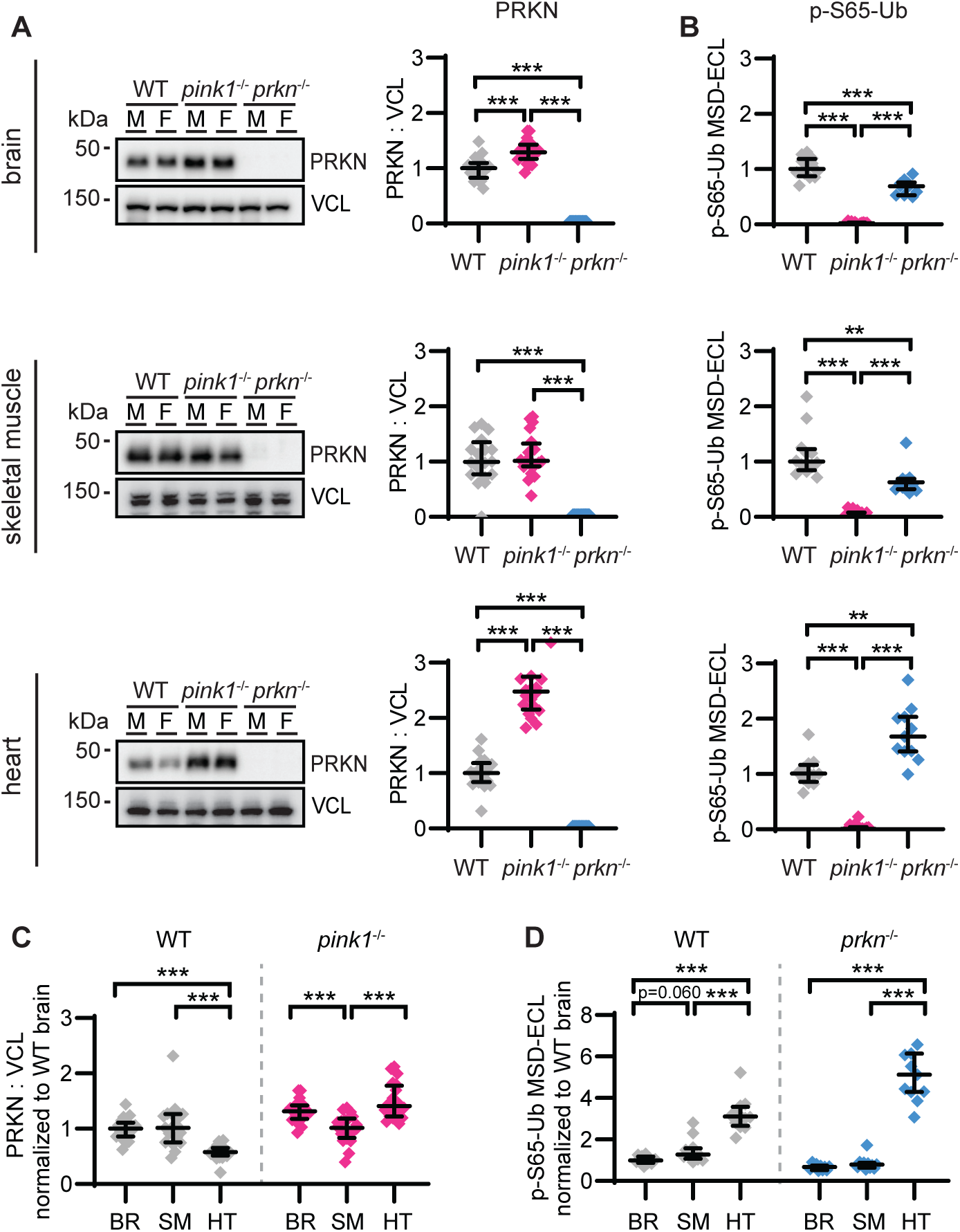
Basal PINK1-PRKN signaling in different tissue from young wild-type and *pink1*^-/-^ or *prkn*^-/-^ mice. PRKN and p-S65-Ub levels were measured in brain, skeletal muscle, and heart from 3-month-old WT, *pink1*^-/-^, and *prkn*^-/-^ mice by western blot or sandwich ELISA, respectively. (**A**) Representative western blots of PRKN with vinculin (VCL) as loading control are shown for the three organs (left panel) and their corresponding densitometric quantifications (right panel). M and F indicate male and female, respectively. To integrate the data from different blots and across different organs, the same two brain samples were loaded on each gel and used as calibrators after densitometric analysis. (**B**) MSD ELISA quantification of p-S65-Ub levels shown as electrochemiluminescence (ECL) signal in the three organs. Data is normalized with WT in the corresponding organs set to 1. (**C**) Quantification of PRKN levels in WT (gray dots) and *pink1*^-/-^ (pink dots) mice across brain (BR), skeletal muscle (SM), and heart (HT) with WT brain set to 1. (**D**) Quantification of p-S65-Ub levels across the same three organs within WT (gray dots) and *prkn*^-/-^ (blue dots) mice with WT brain set to 1. n=12 WT (6M/6F), 20 *pink1^-/-^* (10M/10F), and 11 *prkn^-/-^* (6M/5F). Data are shown as median ± interquartile range. Statistical analysis was performed using the Kruskal-Wallis test and then Mann- Whitney test followed by Bonferroni correction for multiple comparisons (*p<0.05, **p<0.01, ***p<0.001).

As a readout of the joint PINK1 Ub kinase and PRKN Ub ligase activity we measured p-S65-Ub levels using a Meso Scale Discovery (MSD) sandwich ELISA [28]. WT mice had significantly higher p-S65-Ub levels compared to the *pink1*^-/-^ animals, suggesting that the pathway is active under basal conditions in all three organs (**Fig. 1B**). Similar to previous results, p-S65-Ub levels in *prkn*^-/-^ mice compared to WT [29] were reduced by about 30% in brain and to a similar extent in skeletal muscle, reiterating that PRKN amplifies the formation of pS65-Ub in a positive feedback loop. Surprisingly, *prkn*^-/-^ hearts showed strongly elevated p-S65-Ub levels with almost 1.7-fold increase compared to hearts of age-matched WT mice, suggesting that different mechanisms are at play here.

Comparison of PRKN protein levels across the three organs in WT mice revealed similar levels in brain and skeletal muscle at 3-months-old, but much lower levels in heart reaching only 58% in comparison (**Fig. 1C**). Yet, in the absence of PINK1 at this age, PRKN protein levels were relatively similar in brain and heart, but lower in skeletal muscle of *pink1*^-/-^ mice. Comparison of p-S65-Ub across the organs in WT mice revealed slightly increased levels in skeletal muscle (∼1.3-fold) compared to brain but much higher levels in heart (∼3.1-fold) (**Fig. 1D**). In absence of PRKN, this difference in p-S65-Ub was even further pronounced rising to a more than 5-fold p-S65-Ub increase in heart from *prkn*^-/-^ mice compared to the other two organs. Except for slightly enhanced p-S65-Ub levels in *prkn*^-/-^ males compared to females, no other sex differences were observed for the levels of PRKN or p-S65-Ub (**Fig. S1C-D)**.

Taken together, the combined analysis of mRNA, PRKN protein, and especially p- S65-Ub, revealed that the PINK1-PRKN pathway is active in all investigated organs, but to a different extent. Furthermore, data from the heart suggest a distinct regulation of PINK1-PRKN signaling that might be caused by differences in basal activation of flux through the mitophagy pathway.

### Aging increases PINK1-PRKN signaling in brain, skeletal muscle, and heart of mice

We next expanded our analyses of PINK1-PRKN signaling in the three organs to an aging cohort consisting of young (3-month-old), middle-aged (13-month-old), and old (23-month-old) WT mice. Quantitative RT-PCR revealed that *Pink1* mRNA levels were significantly elevated with age in all three organs (1.3- to 1.5-fold) when comparing 3- and 23-month-old mice (**Fig. S2A**). *Prkn* mRNA levels generally increased with age across organs as well (1.2- to 1.6-fold) but were only statistically significant in brain tissue (**Fig. S2B)**.

In contrast to mRNA, PRKN protein levels did not alter with age across all tissue types (**Fig. 2A**). However, p-S65-Ub levels were significantly higher in older mice in all three organs with the largest increase in skeletal muscle (∼2.4-fold), followed by heart (∼1.7-fold) and brain (∼1.5-fold) when comparing the youngest and oldest age groups (**Fig. 2B**). There was no significant correlation between mRNA and protein levels of PRKN in any of the tissues (**Fig. S2C**). Likewise, levels of PRKN protein and the mitophagy label p-S65-Ub did not correlate (**Fig. S2D**). Yet, gene expression of both *Pink1* and *Prkn* strongly correlated with the respective p-S65-Ub levels in all tissues across age (**Fig. 2C- D**). Relative to the brain, PRKN protein amounts were similar in skeletal muscle, but significantly lower in the heart at 3 and 13 months of age (**Fig. 2E**). In contrast, relative to the brain p-S65-Ub levels were greater in skeletal muscle and much higher in the heart at all ages (**Fig. 2F**).

**Figure 2.**
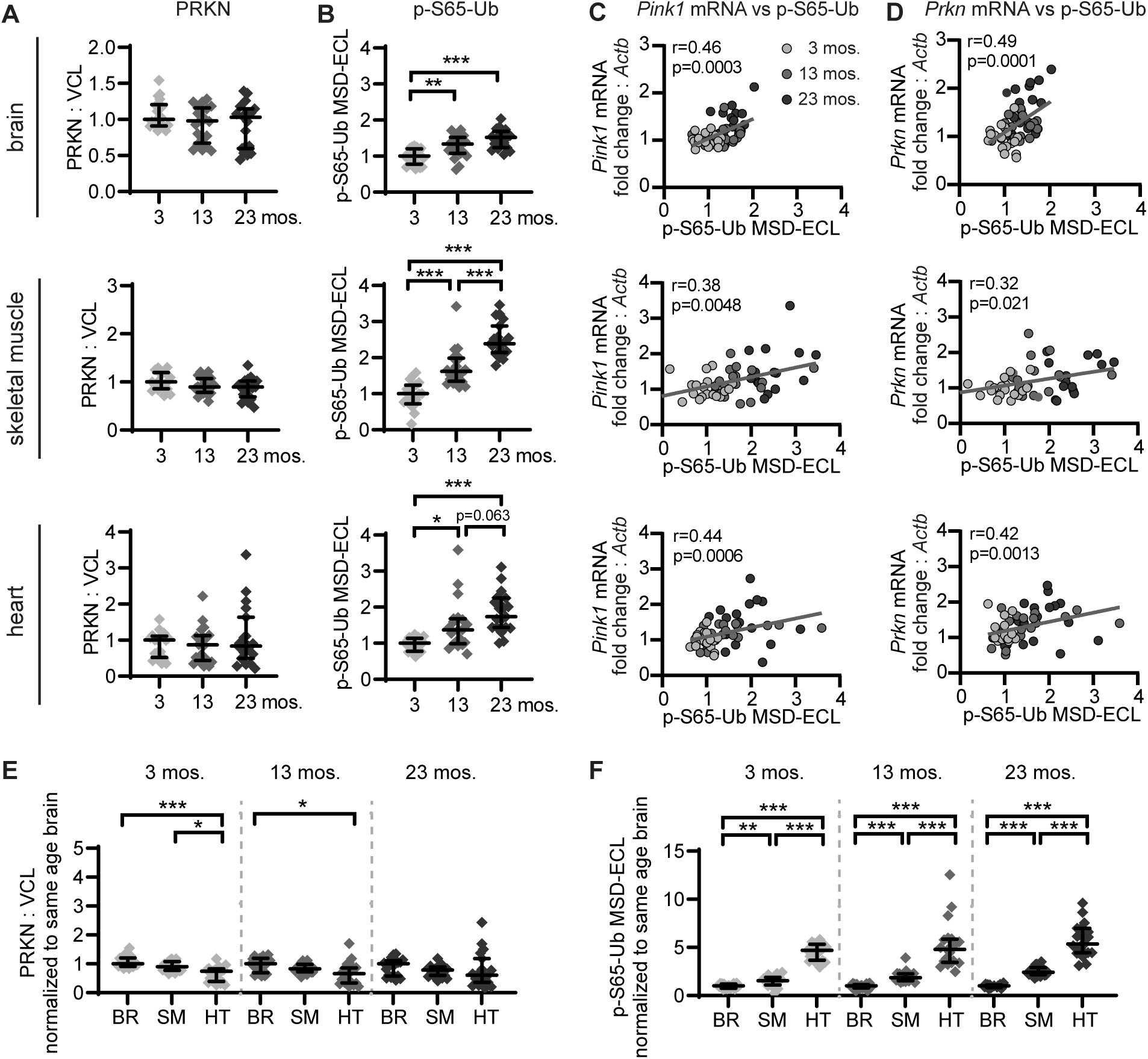
Age-dependent increase of p-S65-Ub and correlation with *Pink1* or *Prkn* expression in different mouse tissue in different mouse tissue. PRKN and p-S65-Ub levels were measured in brain, skeletal muscle, and heart from 3-, 13-, and 23-month-old WT mice by western blot or sandwich ELISA, respectively. (**A**) Densitometric quantifications of PRKN protein levels in western blot relative the loading control VCL. To integrate the data from different blots and across different organs, the same two brain samples were loaded on each gel and used as calibrators after densitometric analysis. (**B**) MSD ELISA quantification of p-S65-Ub levels shown as ECL signal in the three organs during aging. Data is normalized with 3-month-old WT in the corresponding organ set to 1. (**C, D**) Spearman correlation of p-S65-Ub levels with *Pink1* mRNA (**C**), and *Prkn* mRNA (**D**) levels. (**E, F**) Comparison of PRKN (**E**) and p-S65-Ub (**F**) levels across brain (BR), skeletal muscle (SM), and heart (HT) within the same age groups with WT brain in each age group set to 1. n=19-20/age group (10M/10F). Data are shown as median ± interquartile range. Groupwise comparison was performed using the Kruskal-Wallis test and then Mann-Whitney test followed by Bonferroni correction for multiple comparisons (*p<0.05, **p<0.01, ***p<0.001).

We noted a few statistically significant sex differences during aging (**Fig. S3**). In young animals, levels of *Pink1* mRNA were elevated in skeletal muscle of males, while levels of p-S65-Ub were increased in brains of males compared to females. The most pronounced difference was observed in heart with females having much greater p-S65- Ub levels compared to males at mid age. However, even larger samples sizes may be needed to confirm these findings and the biological significance remains to be established in each case.

In summary, during aging we identified increases in *Pink1* and *Prkn* mRNA as well as in p-S65-Ub levels across all tissues. PRKN protein levels however rather declined consistent with the notion that increased activation also facilitates the turnover of the E3 Ub ligase. In this context, it is noteworthy that compared to other tissues, heart showed the least amount of PRKN protein but the highest levels of p-S65-Ub.

### Aging intensifies tissue-specific PINK1-PRKN signaling

Next, we sought to further evaluate findings in *pink1*^-/-^ and *prkn*^-/-^ animals at later ages and using independent methods. Therefore, we utilized different sets of WT and *pink1*^-/-^ or *prkn*^-/-^ mice with young (4-month-old) or advanced age (26-month-old). Consistent with the results from the larger aging cohort (Fig. 2), there was no significant change in PRKN levels in WT animals over age (**Fig. 3A**). The increase of PRKN protein in young *pink1*^-/-^ mice was weaker and statistical significance was lost for brain in this smaller cohort (**Fig. 3A**), compared to Fig. 1. However, PRKN levels were increased 1.6- to 2.6-fold in brain and in skeletal muscle of old *pink1*^-/-^ compared to age-matched WT animals, respectively. In the heart, PRKN levels were significantly increased in young *pink1*^-/-^ mice compared to young WT animals and the fold change between the two genotypes remained similar with age. p-S65-Ub levels increased over age in all tissue types (**Fig. 3B**), although the significance for brain, which showed the smallest fold change of all organs in the larger cohort (Fig. 2), was lost. In young *prkn*^-/-^ mice, p-S65-Ub levels were decreased in brain, but not in skeletal muscle compared to age-matched WT mice (**Fig. 3B**), consistent with Fig. 1. There was no further change with age in either tissue type. In heart, p-S65-Ub levels were again strongly increased in young *prkn*^-/-^ mice (2.1-fold) and further elevated in old *prkn*^-/-^ animals compared to age-matched WT (3.3-fold).

**Figure 3.**
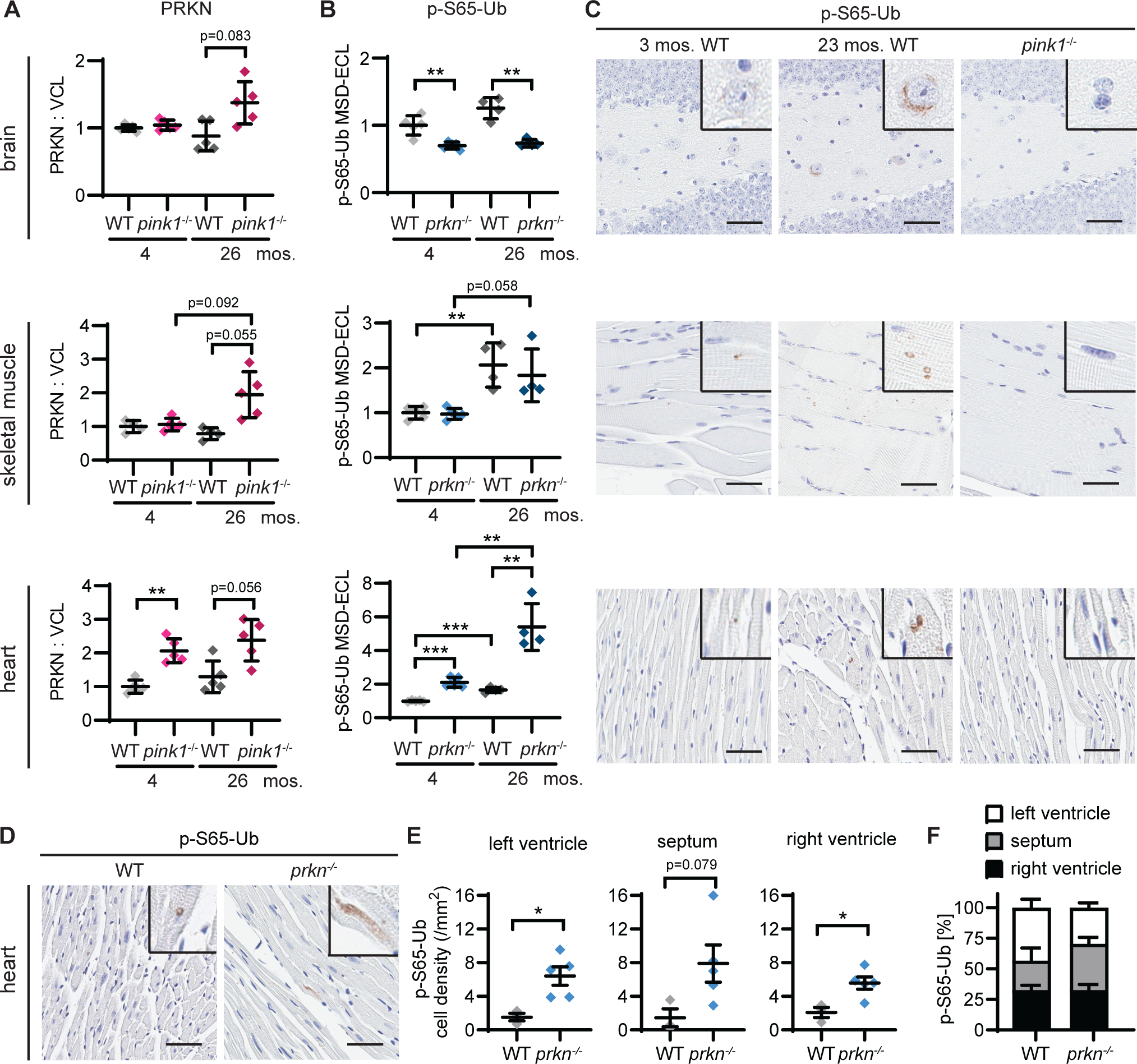
Age-dependent alterations of basal PINK1-PRKN signaling in organs from *pink1^-/-^* and *prkn^-/-^* mice. (**A**, **B**) Brain, skeletal muscle, and heart of young (4-month-old) and old (26-month-old) WT, *pink1*^-/-^ *or prkn^-/-^* mice were analyzed for levels of PRKN (**A**, for WT and *pink1^-^*^/-^) by western blot and of p-S65-Ub (**B**, for WT and *prkn*^-/-^) by sandwich ELISA. Each data is normalized to the 4-month-old WT values set to 1. n=5/genotype/age group. (**C**) Immunohistochemical assessment of p-S65-Ub levels in brain, skeletal muscle, and heart from 3- and 23-month-old WT mice. Corresponding tissues from 14-month-old *pink1*^-/-^ mice were used as negative control. Zoom-in images are shown in black boxes in the top right corner. (**D**) Immunohistochemical staining of p-S65-Ub in heart from 14- month-old WT and *prkn*^-/-^ mice. (**E, F**) p-S65-Ub positive cell density (**E**) and the distribution of positive cells (**F**) were manually quantified in three heart subregions, as indicated. n=3 for WT and n=5 for *prkn^-/-^* mice. Scale bar: 50 µm. Data are shown as mean ± SEM. Statistical analysis was performed using student t-test followed by Bonferroni correction for multiple comparisons (*p<0.05, **p<0.01, ***p<0.001). There was no statistical significance for the comparison between heart regions.

To visualize the results, we examined p-S65-Ub by staining paraffin sections of brain, skeletal muscle, and heart from young and old WT mice. Tissues from *pink1*^-/-^ animals that were used as negative control were devoid of p-S65-Ub signal. In young WT mice, p-S65-Ub immunoreactivity was very faint in all three tissues. Tissue from older WT mice, however, showed overall more frequent and more intense p-S65-Ub signal consistent with the biochemical results (**Fig. 3C**). To further investigate the elevated levels of p-S65-Ub observed in heart upon loss of the Ub ligase PRKN, we stained sections and compared p-S65-Ub levels amongst age-matched 14-month-old WT and *prkn*^-/-^ mice **(Fig. 3D**). Indeed, staining independently showed a significant increase of p-S65-Ub signal in hearts from *prkn*^-/-^ compared to WT controls with an overall 3.7-fold change (**Fig. 3E**), comparable to the 2.1- or 3.3-fold increase measured by MSD in the 4- and 26-month-old cohort, respectively. This signal was increased in both the left and right ventricle with no obvious change in p-S65-Ub distribution between these different compartments when comparing WT and *prkn*^-/-^ mice **(Fig. 3F**).

Taken together, our results confirm and moreover highlight both similarities and distinctions with regard to PRKN protein and p-S65-Ub levels in different tissues with advanced age. Normal aging seems to amplify some of the genotype-specific effects in mitophagy further suggesting distinct activation and turnover rates between different tissues, with heart being the most affected.

## DISCUSSION

PINK1-PRKN mitophagy has been recognized as an important mitochondrial quality control pathway, but this has been extremely difficult to ascertain *in vivo* in mice. Using a biochemical approach, we have previously shown that PINK1-PRKN signaling is active in mouse brain even at baseline [28,29]. Here we extended our prior study and compared brain to other energetically demanding tissues such as skeletal muscle and heart in young animals and during aging. Combined analyses of gene expression, protein levels, and enzymatic activities revealed genotype-specific and age-dependent differences regarding the regulation and extent of PINK1-PRKN signaling in these organs. The current assessment of activities and fluxes at baseline provides a framework for further and comparative investigations of the PINK1-PRKN pathway under various stress and disease relevant conditions.

Our data demonstrated that the PINK1-PRKN pathway is broadly active in mice *in vivo*, even at young age. Brain, skeletal muscle, and heart from WT mice all had significantly higher p-S65-Ub levels compared to the same organ from *pink1*^-/-^ mice. In young mice, p-S65-Ub levels in heart were about 3- to 5-fold higher, while levels in skeletal muscle were about 1.3- to 1.5-fold higher than in brain. The widespread basal activity of the pathway is in line with a generally broad expression of *PINK1* and *PRKN* in different cell and tissue types. Consistent with a general age-dependent increase of p-S65-Ub seen in human autopsy brain [16,20], p-S65-Ub levels also increased with aging in mouse tissues (from 3 to 23 months) with the least increase seen in brain (1.5-fold), followed by heart (1.7-fold), and the most increase seen in skeletal muscle (2.4-fold). In aged animals the difference in PINK1-PRKN signaling between the different organs was even more pronounced. At 23 months, heart and skeletal muscle had 5.4-fold and 2.4-fold higher p- S65-Ub levels compared to brain, respectively. The generation of p-S65-Ub is primarily driven by mitochondrial damage, but a decline in autophagic-lysosomal degradation could also contribute to its build-up. Here, we found a general increase of both *Pink1* and *Prkn* mRNA with aging in all three organs, suggesting age-dependent gene expression changes may contribute to increased PINK1-PRKN signaling.

Recently, we described that endogenous PRKN protein accumulates in the absence of PINK1, indicating that its activation and turnover are tightly linked [29]. Consistently, PRKN levels were slightly but significantly increased in young *pink1^-/-^* brain here as well (1.3-fold) at least in the larger cohort. While we did not observe a statistically significant effect at 3-4 months of age, another recent study reported increased PRKN protein levels in skeletal muscle of 6-month-old *pink1^-/-^* animals [30]. The most robust effect was observed in heart. Here, loss of PINK1 resulted in an almost 2.5-fold increase in PRKN protein. WT hearts displayed the greatest levels of p-S65-Ub but had the lowest PRKN protein relative to the other organs. Together this suggest that basal PINK1-PRKN signaling is most active in heart and this may also be reflected by the cardiac phenotypes of knockout mice that appear most prominent under stress or disease conditions [31–34]. The respective fold increase in PRKN protein seen in *pink1*^-/-^ tissue seemed more pronounced with age. Moreover, in WT animals, PRKN protein levels rather declined with age as the the enzyme may be increasingly activated and “consumed” by mitophagy [29]. However, more animals are certainly needed to substantiate both findings. Collectively, our data suggest the changes observed in relative PRKN protein levels in WT or *pink1*^-/-^ animals might be used as surrogate for distinct basal mitophagy rates across the different tissues and ages.

PRKN is known to engage in a positive feedback loop during which the Ub ligase PRKN provides more substrates for the Ub kinase PINK1 [35–37]. Accordingly, in brain and skeletal muscle from *prkn*^-/-^ animals, the levels of the joint product p-S65-Ub were reduced by more than 1.5-fold. This is similar to what is seen with PRKN-deficient cell cultures in which the pathway has been maximally activated by chemical depolarization of mitochondria [29]. In stark contrast however, p-S65-Ub levels were 1.6-fold increased in *prkn*^-/-^ hearts, pointing towards a mechanistic difference of mitophagy in heart vs the other organs. This effect was even further pronounced in hearts from older *prkn*^-/-^ animals. The heart-specific increase of p-S65-Ub in *prkn*^-/-^ was independently confirmed by immunohistochemistry and was detected rather broadly across heart subregions. The elevated p-S65-Ub levels were not a result of increased *Pink1* expression as transcript levels did not change in absence of PRKN. While the underlying mechanisms remain uncertain, mechanistic explanations include other potential compensatory changes amongst enzymes that may affect de-/conjugation of p-S65-Ub or a block in degradation as suggested in PRKN deficient flies, which also present with increased p-S65-Ub levels [38].

The biochemical assessment of p-S65-Ub is selective for PINK1-PRKN signaling and more sensitive than fluorescent reporters that measure all mitophagy pathways collectively [39–42]. However, we have only been able to quantify protein levels of the E3 Ub ligase PRKN, but not of the Ub kinase PINK1 given to the current inability of the field to reliably detect the mouse protein. Measures of mitochondrial mass and damage as well as autophagic-lysosomal capacity could provide additional context and may inform about the relative activation of and flux through the PINK1-PRKN pathway in each tissue. Beyond the organs described in detail, there was a general increase in p-S65-Ub in other mouse tissue during aging too, but there was also lot more variability noted especially with more advanced age. While some of this is likely due to different organ-specific matrix effects, it is important to note that PINK1-PRKN signaling is a dynamic pathway that cycles between activation and degradation. Thus, some variation can be expected *in vivo* at any time and this might be further influenced by stress due to housing conditions, caloric intake, activity, and/or infections. Overall, even larger samples sizes may be needed, especially at older age, and to fully validate suggestive sex differences. In this context it will be important to determine whether one sex is driving the observed effects in presence or absence of PINK1 or PRKN.

Altogether, our work demonstrates that PINK1-PRKN signaling is constitutively active *in vivo* in mouse. In brain, skeletal muscle, and heart, we found significant differences in regulation and basal activity of the PINK1-PRKN pathway during normal aging. Our analysis complements existing broad mitophagy reporter readouts and sets the stage for follow-up analyses during mtDNA mutagenic stress or exhaustive exercise, and in the context of other genetic or pharmacological challenges known to further activate PINK1-PRKN signaling.

## MATERIALS AND METHODS

### Mice

All mouse procedures were performed in accordance with the Mayo Clinic Institutional Animal Care and Use Committee policies. *pink1*^-/-^ and *prkn*^-/-^ mice (Jackson Laboratories, strains 017946 and 006582, respectively) [43,44] were backcrossed over 3 generations to a C57BL/6J background (Jackson Laboratories, 00664). The final cohort to study genotype differences consisted of 3-month-old ± 2 weeks WT (n=12, 6 males and 6 females), *pink1*^-/-^ (n=20, 10 males and 10 females), and *prkn*^-/-^ mice (n=11, 6 males and 5 females). A cohort of WT mice (C57BL/6J) of different ages was received from the National Institute on Aging (NIA). The cohort consisted of 59 mice: 3-month-old ± 2 weeks (n=20, 10 males and 10 females), 13-month-old ± 2 weeks (n=20, 10 males and 10 females) and 23-month-old ± 2 weeks (n=19, 10 males and 9 females).

Additional cohorts were used to study genotype effects during aging and for further validation. For WT and *pink1*^-/-^ the cohort consisted of 20 mice: young (4-month-old) WT (n=5, 3 males and 2 females) and *pink1*^-/-^ (n=5, 3 male and 2 females), and old (26-month-old) WT (n=5, 3 males and 2 females) and *pink1*^-/-^ (n=5, 3 male and 2 females). For WT and *prkn*^-/-^, we received tissue from Dr. Nobutaka Hattori (Juntendo University, Japan) [45]. The tissue was harvested from young (4-month-old ± 2 weeks) WT (n=5, all males), and *prkn*^-/-^ (n=5, all males), and old (26-month-old ± 2 weeks) WT (n=4, all males) and *prkn*^-/-^ animals (n=4, all males). For imaging analysis, two young (3-month-old, 1 male and 1 female) and old (23-month-old, 1 male and 1 female) WT mice were used for immunohistochemical staining across different organs. Tissues from two *pink1*^-/-^ mice (14- month-old, 1 male and 1 female) were used as negative control. Additionally, a group of 14-month-old ± 2 weeks WT (n=3, all males) and *prkn*^-/-^ (n=5, 3 males and 2 females) was used for immunohistochemical analysis in heart.

### Mouse tissue

All mice were anesthetized and perfused intracardially with cold PBS (Thermo Fisher Scientific, 14190235). Brain, skeletal muscle, and heart were harvested from each mouse and immediately flash frozen using liquid nitrogen. Frozen tissues were stored at −80 °C and remained frozen on dry ice during weighing. Mouse tissues were homogenized using 2 mL glass homogenizers (Fisher Scientific, K885300-0002) in 5x volumes of ice-cold TBS buffer (50 mM Tris pH 7.4 [Thermo Scientific, J75825-A4], 150 mM NaCl [Fisher Scientific, BP358-10]) containing phosphatase and protease inhibitors (Roche, 04906845001, 05056489001) (TBS+). Both plunger A and then B, were used for 15 strokes each and TBS+ homogenates aliquoted and flash frozen in liquid nitrogen until further extraction.

### RNA extraction and quantitative RT-PCR

A total of 50 µL TBS homogenate from mouse brain, skeletal muscle, and heart was used for RNA extraction using a RNeasy Plus Mini Kit (Qiagen, 74134). Quantitative RT-PCR was carried out using iTaq Universal SYBR Green One-Step Kit (Bio-Rad, #1725150). 50 ng of RNA was mixed with primers for targeted genes, SYBR Green, and iScript reverse transcriptase in a 5 µL reaction. The PCR was executed using a 384-well block on a LightCycler 480 system (Roche, Switzerland). Relative transcript levels for *Pink1* and *Prkn* were calculated with 2-^ΔΔCT^ [46] using *Actb* as housekeeping gene and normalized to the relative expression level of the corresponding tissue from 3-month-old mice. The following primers were used: *Pink1* forward 5’-GAGTGGGACTCAGATGGCTGTCC-3’, *Pink1* reverse 5’-CCAGAATGGGCTGTGGACACCTC-3’, *Prkn* forward 5’- GATTCAGAAGCAGCCAGAGG-3’, *Prkn* reverse 5’-GGTGCCACACTGAACTCG-3’, *Actb* forward 5’-AGTGTGACGTTGACATCCGTA-3’, *Actb* reverse 5’- GCCAGAGCAGTAATCTCCTTC-3’.

### Protein extraction

Tissue lysis and protein extraction was completed by adding 25 µL of 5x RIPA buffer (50 mM Tris, pH 8.0 [Thermo Scientific, J75825-A4], 150 mM NaCl [Fisher Scientific, BP358- 10], 0.1% SDS [Fisher Bioreagents, BP166-500], 0.5% deoxycholate [Sigma, D6750], 1% NP-40 [United States Biochemical, 19628]) to 100 µL TBS homogenate, vortexed and incubated for 30 minutes on ice. During the tissue homogenization and lysing steps, homogenates and lysates were always placed on ice. After incubation, lysates were centrifuged twice at 20,000 x *g* for 15 minutes at 4 °C to remove insoluble components, lipids, and nucleic acids. The resulting supernatant was transferred to a fresh tube each time. Protein concentration was determined using a Bicinchoninic acid assay (ThermoFisher, 23225).

### Western blot

Protein lysates were prepared in Laemmli buffer (62.5 mM Tris HCl pH 6.8, 1.5% SDS [w:v], 8.33% glycerol [v:v] [Fisher Scientific, BP229-1], 1.5% β-mercaptoethanol [v:v] [Sigma, M3148], 0.005% bromophenol blue [w:v] [Sigma, 5525]) and boiled at 95°C for five minutes. Protein electrophoresis was performed using 8-16% Tris-Glycine gels (Invitrogen, XP08165BOX) and standard running buffer (25 mM Tris, 0.2 M Glycine, 0.1% SDS) at room temperature. 30 µg of total protein was loaded per well. For comparison between blots and across organs we loaded the same two brain samples on each gel and used them as calibrators after densitometric analysis. Proteins were transferred onto polyvinylidene fluoride (PDVF) membranes (Millipore, IEVH00005). Blocking was performed with 5% non-fat milk dissolved in TBST for 1 hour followed by incubation with primary antibodies PRKN (Cell Signaling Technology, 2132) at 1:2000 dilution in 5% BSA in TBST at 4 °C overnight and VCL (Sigma-Aldrich, V9131) as the loading control at 1:375,000 dilution in 5% milk for 1 hour at room temperature. Next, the membranes were washed for 30 minutes in TBST and incubated with the secondary HRP-conjugated antibody (Jackson Immuno research, 111-035-003 [anti-rabbit], 715-035-150 [anti-mouse]) in 5% milk for 1 hour at room temperature. The signal was developed using an HRP Substrate (Millipore, WBKLS0500) and imaged using a ChemiDoc MP imaging system (Bio-Rad, Hercules, CA, USA). Western blot signal intensity was determined using the ImageStudio Lite software (version 5.2).

### P-S65-Ub Meso-Scale Discovery sandwich ELISA

Our assay to detect p-S65-Ub has been described before [28]. Briefly, the 96-well MSD plates (Meso Scale Diagnostics, L15XA) were coated with 1 µg/mL p-S65-Ub capturing antibody (Cell Signaling Technology, 62802), blocked with 1% BSA in TBST. Then 30 µg/30 µl protein lysates were added to each well and after 2 h incubation, 5 µg/mL ubiquitin antibody (Thermo Fisher Scientific, 14-6078-37) was used for detection followed by a sulfo-tag coupled secondary antibody (Meso Scale Diagnostics, R32AC-1). The plates were then read on a MESO QuickPlex SQ 120 reader (Meso Scale Diagnostics, Rockville, MD, USA). All samples were run in duplicates and each plate had an organ specific *pink1*^-/-^ sample included as a negative control as well as a recombinant K48 p-S65-Ub tetramer (Boston Biochem, UC-250) as a positive control. The raw values were background corrected by subtracting the organ-specific *Pink1^-/-^* value.

### Immunohistochemistry

Immunohistochemical staining was performed with paraffin-embedded mouse sagittal brain, skeletal muscle, and ventricular heart sections (one section per organ for each animal). Sections were cut at a thickness of 5 microns and mounted onto positively charged slides to dry overnight at 60°C. After sections were deparaffinized and rehydrated, antigen retrieval was performed by steaming the sections for 30 minutes in deionized water. Immunostaining was carried out using Envision Plus kit (Agilent, K4011). Endogenous peroxidase blocking was performed for 5 minutes using hydrogen peroxide (Swan, L0011380FB). Following blocking with 5% normal goat serum (Invitrogen, 16210- 072) for 20 minutes, sections were incubated with primary antibody against pS65-Ub (in-house, 1:250) [20] that was diluted in Dako Antibody Diluent with Background Reducing Components (Agilent, S3022) for 45 minutes at room temperature. Subsequently, sections were incubated with EnVision-Plus anti-rabbit labeled polymer-HRP (Agilent, K4003) for 30 minutes at room temperature. Peroxidase labeling was visualized with the chromogen solution 3,3’-diaminobenzidine (Agilent, K3468). Sections were then counterstained with Gill 1 hematoxylin (Epredia, 6765006) and following dehydration, coverslipped with Cytoseal XYL mounting media (Epredia, 8312-4). After drying, sections were scanned with an Aperio AT2 digital pathology scanner (Leica Biosystems, Wetzlar, Germany) at 20x magnification.

### Statistical analysis

Data were analyzed using GraphPad Prism (version 9.2.). Given that most measures were not normally distributed and had differing variances between groups, non-parametric tests (Kruskal-Wallis and Mann-Whitney U tests followed by adjustment with Bonferroni correction) were used. For groups with sample size less than 6, a parametric test (Student’s t-test followed by adjustment with Bonferroni correction) were used. Correlation analysis was performed using Spearman’s test. For Bonferroni correction, all tests were corrected for three organs and additionally for multiple pair-wise comparisons between age, genotype, and/or sex groups.

## ACKNOWLEDGEMENTS

We thank the Neuropathology laboratory at Mayo Clinic Florida under the leadership of Dr. Dennis Dickson for processing and immunohistochemical staining of all mouse tissues. We also thank the National Institute on Aging (NIA) for providing young and aged WT mice.

## FUNDING

This work was supported in part by a grant from the Michael J. Fox Foundation for Parkinson’s Research (MJFF-019046 to I.G.G., M.S.G., K.L.S., and W.S.). Work in the authors labs is further funded by the National Institutes of Health (R56 AG062556, RF1 NS085070, R01 NS110085, and U54 NS110435 to W.S.), the Department of Defense Congressionally Directed Medical Research Programs (CDMRP) (W81XWH-17-1-0248 to W.S.), the Florida Department of Health - Ed and Ethel Moore Alzheimer’s Disease Research Program (9AZ10 to W.S. and 22A07 to F.C.F.), the Ted Nash Long Life Foundation (W.S.) and The Michael J. Fox Foundation for Parkinson’s Research (W.S. and F.C.F.). W.S. is also supported by Mayo Clinic Foundation, Mayo Clinic Center for Biomedical Discovery (CBD), and Mayo Clinic Robert and Arlene Kogod Center on Aging. X.H. is supported by Mayo Clinic Alzheimer Disease Research Center (ADRC, P30 AG062677) pilot and developmental project grants and research fellowships from the American Parkinson Disease Association (APDA) and the Alzheimer’s Association (AARF-22-973152). Mayo Clinic Florida is an American Parkinson Disease Association (APDA) Center for Advanced Research (W.S., F.C.F., and D.W.D.). The work was supported in part by National Heart Lung and Blood Institute (NHLBI) under award number R01 HL164520 (D.F.). I.G.G. is also funded by the Medical Research Council, UK (MC_UU_00018/2).

## DISCLOSURE STATEMENT

Mayo Clinic, F.C.F., and W.S. hold a patent related to PRKN activators (Small Molecule Activators of Parkin Enzyme Function, US patent, 11401255B2; August 02, 2022). Additional funding sources to disclose but not pertinent to the current study include a grant from Amazentis SA (to W.S.). All other authors declare they have no competing interests. This research was conducted in compliance with Mayo Clinic conflict of interest policies.

## ABBREVIATIONS

ECL: electrochemiluminescence
ELISA: enzyme-linked immunosorbent assay
MSD: Meso Scale Discovery
PD: Parkinson disease
p-S65-Ub: Serine-65 phosphorylated ubiquitin
RT-PCR: real-time polymerase chain reaction
Ub: ubiquitin
WT: wild-type.

## REFERENCES

1. Truban D, Hou X, Caulfield TR, et al. PINK1, Parkin, and Mitochondrial Quality Control: What can we Learn about Parkinson’s Disease Pathobiology? J Parkinsons Dis. 2017;7(1):13–29.

2. Kane LA, Lazarou M, Fogel AI, et al. PINK1 phosphorylates ubiquitin to activate Parkin E3 ubiquitin ligase activity. J Cell Biol. 2014 Apr 28;205(2):143–53.

3. Kazlauskaite A, Kondapalli C, Gourlay R, et al. Parkin is activated by PINK1-dependent phosphorylation of ubiquitin at Ser65. Biochem J. 2014 May 15;460(1):127–39.

4. Koyano F, Okatsu K, Kosako H, et al. Ubiquitin is phosphorylated by PINK1 to activate parkin. Nature. 2014 Jun 5;510(7503):162–6.

5. Okatsu K, Koyano F, Kimura M, et al. Phosphorylated ubiquitin chain is the genuine Parkin receptor. J Cell Biol. 2015 Apr 13;209(1):111–28.

6. Sauve V, Lilov A, Seirafi M, et al. A Ubl/ubiquitin switch in the activation of Parkin. EMBO J. 2015 Oct 14;34(20):2492–505.

7. Wauer T, Simicek M, Schubert A, et al. Mechanism of phospho-ubiquitin-induced PARKIN activation. Nature. 2015 Aug 20;524(7565):370–4.

8. Iguchi M, Kujuro Y, Okatsu K, et al. Parkin-catalyzed ubiquitin-ester transfer is triggered by PINK1-dependent phosphorylation. J Biol Chem. 2013 Jul 26;288(30):22019–32.

9. Kondapalli C, Kazlauskaite A, Zhang N, et al. PINK1 is activated by mitochondrial membrane potential depolarization and stimulates Parkin E3 ligase activity by phosphorylating Serine 65. Open Biol. 2012 May;2(5):120080.

10. Shiba-Fukushima K, Imai Y, Yoshida S, et al. PINK1-mediated phosphorylation of the Parkin ubiquitin-like domain primes mitochondrial translocation of Parkin and regulates mitophagy. Sci Rep. 2012;2:1002.

11. Zhang C, Lee S, Peng Y, et al. PINK1 triggers autocatalytic activation of Parkin to specify cell fate decisions. Curr Biol. 2014 Aug 18;24(16):1854–65.

12. Kitada T, Asakawa S, Hattori N, et al. Mutations in the parkin gene cause autosomal recessive juvenile parkinsonism. Nature. 1998 Apr 9;392(6676):605–8.

13. Valente EM, Abou-Sleiman PM, Caputo V, et al. Hereditary early-onset Parkinson’s disease caused by mutations in PINK1. Science. 2004 May 21;304(5674):1158–60.

14. Pickrell AM, Youle RJ. The roles of PINK1, parkin, and mitochondrial fidelity in Parkinson’s disease. Neuron. 2015 Jan 21;85(2):257–73.

15. Vizziello M, Borellini L, Franco G, et al. Disruption of Mitochondrial Homeostasis: The Role of PINK1 in Parkinson’s Disease. Cells. 2021 Nov 4;10(11).

16. Hou X, Fiesel FC, Truban D, et al. Age- and disease-dependent increase of the mitophagy marker phospho-ubiquitin in normal aging and Lewy body disease. Autophagy. 2018;14(8):1404–1418.

17. Hou X, Watzlawik JO, Cook C, et al. Mitophagy alterations in Alzheimer’s disease are associated with granulovacuolar degeneration and early tau pathology. Alzheimers Dement. 2020 Oct 8;17(3):417–30.

18. Narendra DP. Managing risky assets - mitophagy in vivo. J Cell Sci. 2021 Oct 1;134(19).

19. Wu Y, Jiang T, Hua J, et al. PINK1/Parkin-mediated mitophagy in cardiovascular disease: From pathogenesis to novel therapy. Int J Cardiol. 2022 Aug 15;361:61–69.

20. Fiesel FC, Ando M, Hudec R, et al. (Patho-)physiological relevance of PINK1- dependent ubiquitin phosphorylation. EMBO Rep. 2015 Sep;16(9):1114–30.

21. Leduc-Gaudet JP, Reynaud O, Hussain SN, et al. Parkin overexpression protects from ageing-related loss of muscle mass and strength. J Physiol. 2019 Apr;597(7):1975–1991.

22. Han R, Liu Y, Li S, et al. PINK1-PRKN mediated mitophagy: differences between in vitro and in vivo models. Autophagy. 2022 Nov 3:1–10.

23. Paul S, Pickrell AM. Hidden phenotypes of PINK1/Parkin knockout mice. Biochim Biophys Acta Gen Subj. 2021 Jun;1865(6):129871.

24. Matheoud D, Cannon T, Voisin A, et al. Intestinal infection triggers Parkinson’s disease-like symptoms in Pink1(-/-) mice. Nature. 2019 Jul;571(7766):565–569.

25. Sliter DA, Martinez J, Hao L, et al. Parkin and PINK1 mitigate STING-induced inflammation. Nature. 2018 Sep;561(7722):258–262.

26. Noda S, Sato S, Fukuda T, et al. Loss of Parkin contributes to mitochondrial turnover and dopaminergic neuronal loss in aged mice. Neurobiol Dis. 2020 Mar;136:104717.

27. Hou X, Chen TH, Koga S, et al. Alpha-synuclein-associated changes in PINK1-PRKN-mediated mitophagy are disease context dependent. Brain Pathol. 2023 Sep;33(5):e13175.

28. Watzlawik JO, Hou X, Fricova D, et al. Sensitive ELISA-based detection method for the mitophagy marker p-S65-Ub in human cells, autopsy brain, and blood samples. Autophagy. 2021 Sep;17(9):2613–2628.

29. Watzlawik JO, Fiesel FC, Fiorino G, et al. Basal activity of PINK1 and PRKN in cell models and rodent brain. Autophagy. 2023 Dec 2:1–12.

30. Singh F, Wilhelm L, Prescott AR, et al. PINK1 regulated mitophagy is evident in skeletal muscles. Autophagy Reports. 2024 2024/12/31;3(1):2326402.

31. Kubli DA, Zhang X, Lee Y, et al. Parkin protein deficiency exacerbates cardiac injury and reduces survival following myocardial infarction. J Biol Chem. 2013 Jan 11;288(2):915–26.

32. Gong G, Song M, Csordas G, et al. Parkin-mediated mitophagy directs perinatal cardiac metabolic maturation in mice. Science. 2015 Dec 4;350(6265):aad2459.

33. Dorn GW, 2nd. Parkin-dependent mitophagy in the heart. J Mol Cell Cardiol. 2016 Jun;95:42–9.

34. Tong M, Saito T, Zhai P, et al. Mitophagy Is Essential for Maintaining Cardiac Function During High Fat Diet-Induced Diabetic Cardiomyopathy. Circ Res. 2019 Apr 26;124(9):1360–1371.

35. Lazarou M, Narendra DP, Jin SM, et al. PINK1 drives Parkin self-association and HECT-like E3 activity upstream of mitochondrial binding. J Cell Biol. 2013 Jan 21;200(2):163–72.

36. Ordureau A, Sarraf SA, Duda DM, et al. Quantitative proteomics reveal a feedforward mechanism for mitochondrial PARKIN translocation and ubiquitin chain synthesis. Mol Cell. 2014 Nov 6;56(3):360–375.

37. Shiba-Fukushima K, Arano T, Matsumoto G, et al. Phosphorylation of mitochondrial polyubiquitin by PINK1 promotes Parkin mitochondrial tethering. PLoS Genet. 2014 Dec;10(12):e1004861.

38. Usher JL, Sanchez-Martinez A, Terriente-Felix A, et al. Parkin drives pS65-Ub turnover independently of canonical autophagy in Drosophila. EMBO Rep. 2022 Dec 6;23(12):e53552.

39. McWilliams TG, Prescott AR, Montava-Garriga L, et al. Basal Mitophagy Occurs Independently of PINK1 in Mouse Tissues of High Metabolic Demand. Cell Metab. 2018 Feb 6;27(2):439–449 e5.

40. Liu YT, Sliter DA, Shammas MK, et al. Mt-Keima detects PINK1-PRKN mitophagy in vivo with greater sensitivity than mito-QC. Autophagy. 2021 Nov;17(11):3753–3762.

41. Ganley IG, Whitworth AJ, McWilliams TG. Comment on “mt-Keima detects PINK1-PRKN mitophagy in vivo with greater sensitivity than mito-QC”. Autophagy. 2021 Dec;17(12):4477–4479.

42. Liu YT, Sliter DA, Shammas MK, et al. Comment on “mt-Keima detects PINK1-PRKN mitophagy in vivo with greater sensitivity than mito-QC”. Autophagy. 2021 Dec;17(12):4484–4485.

43. Goldberg MS, Fleming SM, Palacino JJ, et al. Parkin-deficient mice exhibit nigrostriatal deficits but not loss of dopaminergic neurons. J Biol Chem. 2003 Oct 31;278(44):43628–35.

44. Kitada T, Pisani A, Porter DR, et al. Impaired dopamine release and synaptic plasticity in the striatum of PINK1-deficient mice. Proc Natl Acad Sci U S A. 2007 Jul 3;104(27):11441–6.

45. Sato S, Chiba T, Nishiyama S, et al. Decline of striatal dopamine release in parkin-deficient mice shown by ex vivo autoradiography. J Neurosci Res. 2006 Nov 1;84(6):1350–7.

46. Livak KJ, Schmittgen TD. Analysis of relative gene expression data using real-time quantitative PCR and the 2(-Delta Delta C(T)) Method. Methods. 2001 Dec;25(4):402–8.

