## Supplemental Materials for "Alterations of PINK1-PRKN signaling in mice during normal aging"

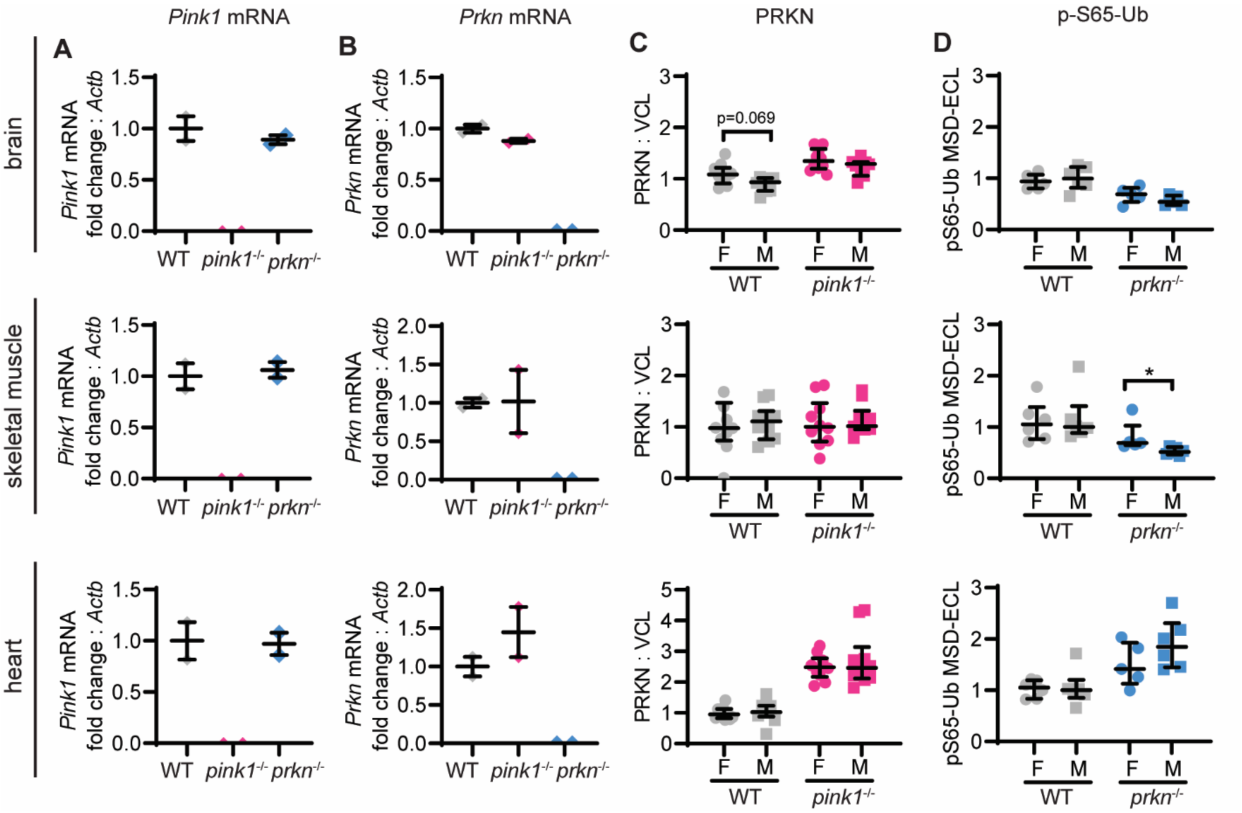


**Supplementary Figure 1. Gene expressions and sex differences in basal PINK1-PRKN signaling in young *pink1*$^{-/-}$ and *prkn*$^{-/-}$ mice.** (**A**, **B**) *Pink1* and *Prkn* mRNA levels were measured in brain, skeletal muscle, and heart from 3-month-old wildtype (WT), *pink1*$^{-/-}$*,* and *prkn*$^{-/-}$ mice by realtime qPCR. *Actin* was used as housekeeping gene. *Pink1* (**A**) and *Prkn* (**B**) mRNA levels were compared between genotypes in each tissue. n=2/genotype. (**C, D**) PRKN and p-S65-Ub protein levels were measured in the same tissue by western blot (for PRKN) or sandwich ELISA (for p-S65-Ub) and were compared between female and male mice within each genotype. (**C**) Densitometric quantifications of PRKN protein levels in western blot relative the loading control VCL. (**D**) MSD ELISA quantification of p-S65-Ub levels shown as electrochemiluminescence (ECL) signal. Data is normalized with WT in the corresponding organs set to 1 and shown as median ± interquartile range. n=12 WT (6M/6F), 20 *pink1*$^{-/-}$ (10M/10F), and 11 *prkn*$^{-/-}$ (6M/5F). Statistical analysis was performed using the Mann-Whitney test followed by Bonferroni correction for multiple comparisons (*p<0.05).

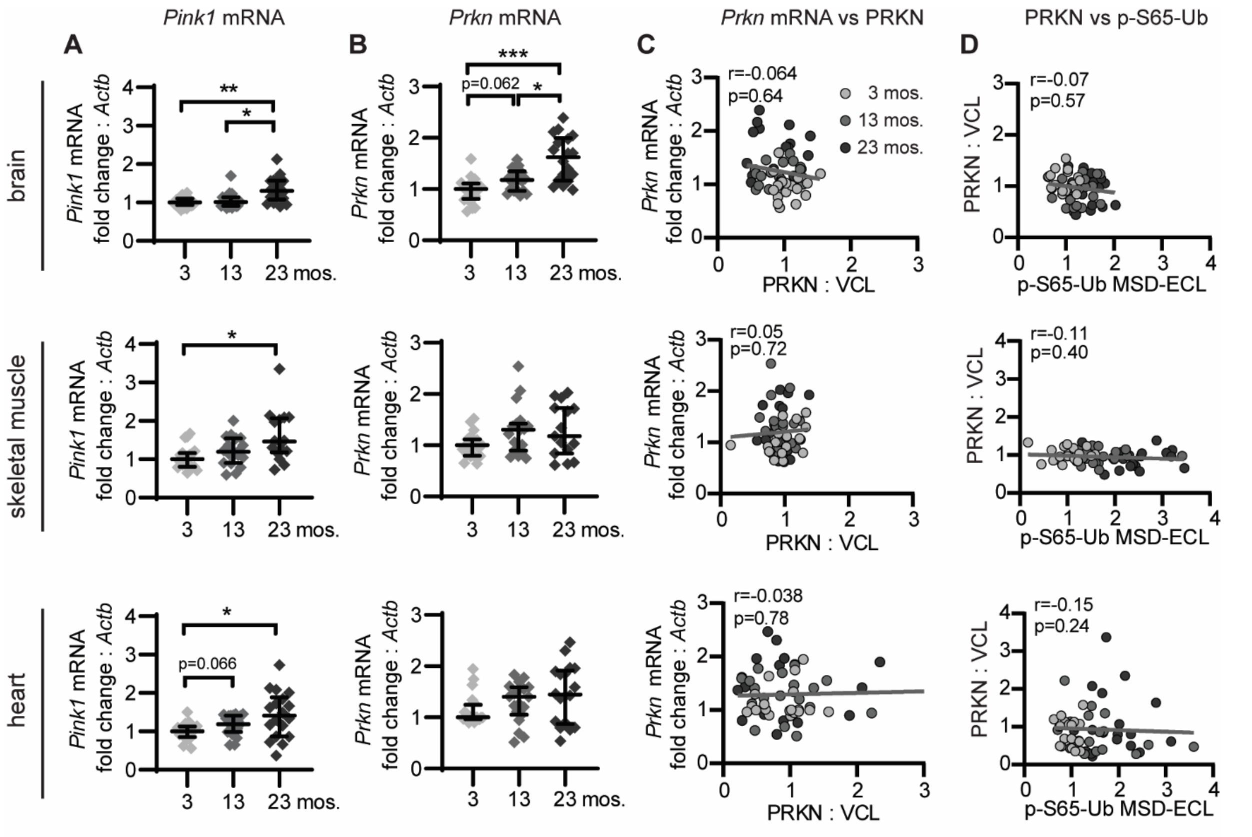


**Supplementary Figure 2. Gene expressions and correlation analysis in basal PINK1-PRKN signaling in mice during aging.** (**A, B**) *Pink1* and *Prkn* mRNA levels were measured in brain, skeletal muscle, and heart from 3-, 13-, and 23-month-old WT mice by realtime qPCR. *Actin* was used as housekeeping gene. *Pink1* (**A**) and *Prkn* (**B**) mRNA levels were compared during aging in each tissue. Data is normalized with 3-month-old WT in the corresponding organ set to 1 and shown as median ± interquartile range. (**C, D**) Correlation of PRKN protein levels with *Prkn* mRNA (**C**), and p-S65-Ub (**D**) levels in the same cohort. n=19-20/age group (10M/10F). Groupwise comparison was performed using Mann-Whitney test followed by Bonferroni correction for multiple comparisons (*p<0.05, **p<0.01, ***p<0.001). Correlation analysis was performed using Spearman's test.

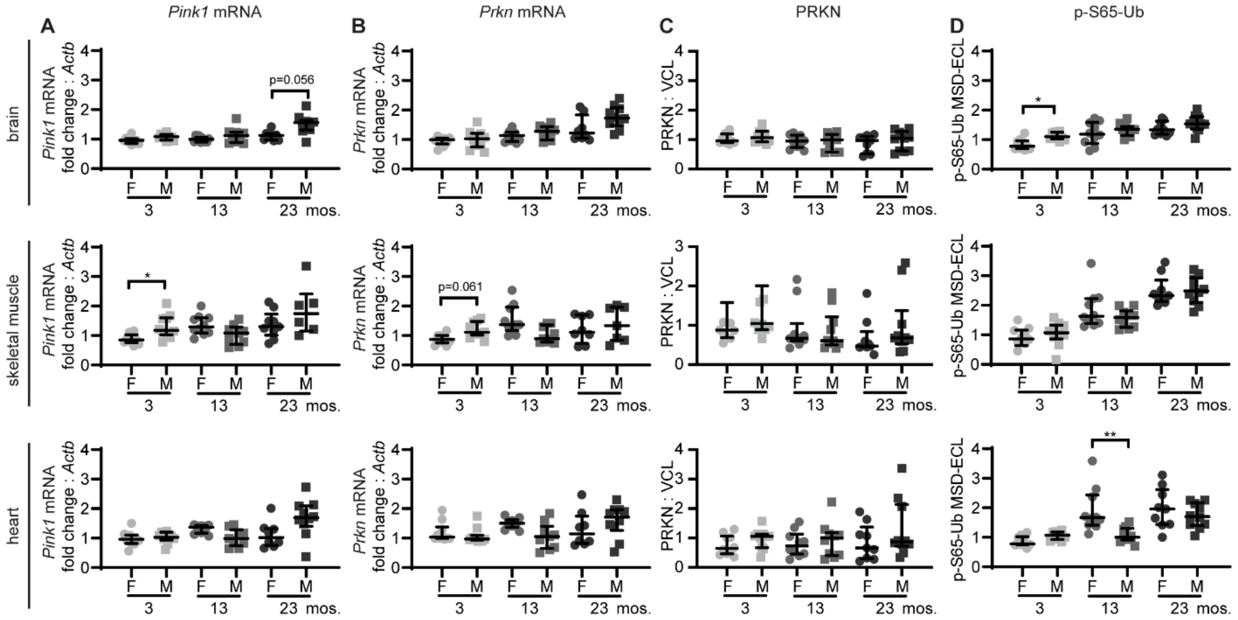


**Supplementary Figure 3. Sex differences in basal PINK1-PRKN signaling in mice during aging.** (**A**, **B**) *Pink1* and *Prkn* mRNA levels were measured in brain, skeletal muscle, and heart from 3-, 13-, and 23-month-old WT mice by realtime qPCR. *Actin* was used as housekeeping gene. *Pink1* (**A**) and *Prkn* (**B**) mRNA levels were compared between female and male mice within each age group. (**C, D**) PRKN and p-S65-Ub protein levels were measured in the same tissue by western blot (for PRKN) or sandwich ELISA (for p-S65-Ub) and were compared between female and male mice within each age group. (**C**) Densitometric quantifications of PRKN protein levels in western blot relative the loading control VCL. (**D**) MSD ELISA quantification of p-S65-Ub levels shown as electrochemiluminescence (ECL) signal. Data is normalized with 3-month-old WT in the corresponding organs set to 1 and shown as median ± interquartile range. n=19-20/age group (10M/10F). Statistical analysis was performed using the Mann-Whitney test followed by Bonferroni correction for multiple comparisons (*p<0.05, **p<0.01).
